# Mechanism of autonomous synchronization of the circadian KaiABC rhythm

**DOI:** 10.1101/2020.11.24.395566

**Authors:** Masaki Sasai

## Abstract

The cyanobacterial circadian clock can be reconstituted by mixing three proteins, KaiA, KaiB, and KaiC, in vitro. In this protein mixture, oscillations of the phosphorylation level of KaiC molecules are synchronized to show the coherent oscillations of the ensemble of many molecules. However, the mechanism of this synchronization remains elusive. In this paper, we explain a theoretical model that considers the multifold feedback relations among the structure and reactions of KaiC. The simulated KaiC hexamers show stochastic switch-like transitions at the level of single molecules, which are synchronized in the ensemble through the sequestration of KaiA into the KaiC-KaiB-KaiA complexes. The proposed mechanism quantitatively reproduces the synchronization that was observed by mixing two oscillating solutions in different phases. The model results suggest that biochemical assays with varying concentrations of KaiA or KaiB can be used to test this hypothesis.

## Introduction

When three cyanobacterial proteins, KaiA, KaiB, and KaiC, are mixed and incubated with ATP in vitro, the phosphorylation level of KaiC shows robust oscillations with approximately 24 h period^1^. Much attention has been focused on this prototypical circadian oscillator, and important features of the oscillations have been elucidated both at the molecule level and in ensemble of molecules^2,3^. At the molecular level, KaiC forms a hexamer^4–6^ and the coordinated binding/unbinding of KaiA and KaiB to/from KaiC generates the phosphorylation rhythm of individual KaiC hexamers^7–12^. At the ensemble level, the autonomous synchronization of a large number of KaiC hexamers produces macroscopic oscillations in a test tube^13,14^. This paper explains a theoretical model that describes the relationship among the structural changes and reactions in individual KaiC molecules, and we discuss how the model explains the experimental data on the ensemble-level synchronization.

Various hypotheses have been proposed to explain the observed synchronization of KaiC hexamers^15–31^. One of the earliest theories was based on the assumption that synchronization is realized through the direct association of multiple KaiC hexamers^15^. However, no experimental evidence has been shown for such assembly of KaiC hexamers in an oscillating solution. Another hypothesis focuses on the assumption of communication among KaiC hexamers through the exchange of KaiC monomers^16–20^. It was argued that this monomer-exchange hypothesis is consistent with the experimental observation on the entrainment of oscillations^13,32^. A third hypothesis was based on the assumption that KaiA may preferentially bind to particular states or complexes of KaiC, which sequestrates KaiA^21–31^. When the total amount of KaiA is limited, this preferential binding of KaiA in the specific states or complexes should deplete free unbound KaiA, resulting in decrease of the binding frequency of KaiA to KaiC in other states. Because binding of KaiA is necessary for promoting phosphorylation of KaiC^7^, a reduction in KaiA binding frequency leads to ‘kinetic congestion’ in the phosphorylation process. Then, an accumulation of the population of KaiC at the specific levels of phosphorylation gives rise to coherent synchronized oscillations. The experimental observation that supports this KaiA sequestration hypothesis is that the ensemble’s oscillations disappear when KaiA is too abundant in the solution^33^. However, the precise mechanism of KaiA sequestration is still not defined. Different scenarios have been proposed with various assumptions. In one, KaiA could be sequestrated into the lowly phosphorylated states in the phosphorylation process^21–25^ or in the dephosphorylation process^26,27^ of KaiC. Another scenario is that KaiA could be sequestrated into the KaiC-KaiB-KaiA complexes that appear during the dephosphorylation process^28–31^. To examine the validity of the KaiA sequestration hypothesis, further quantitative comparisons between theoretical models and experimental data are necessary.

A direct test on the synchronization mechanism is to mix two solutions oscillating in different phases. Ito et al.^13^ showed that when the oscillating solution in the phosphorylation phase and the equal amount of the same solution in the dephosphorylation phase are mixed, the mixture solution’s oscillation is entrained into the dephosphorylation phase. Because the monomer exchange is more frequent in the dephosphorylation phase than in the phosphorylation phase^34^, Ito et al. argued that the monomer exchange could be the mechanism of this entrainment, and that the monomer exchange could explain the autonomous synchronization in stable oscillations^13^. Indeed, a theoretical model assuming the monomer-exchange mechanism reproduced the data of Ito et al^19^. However, careful examination is necessary on whether the KaiA sequestration mechanism is consistent with Ito et al.’s experimental data or is inadequate to explain this phenomenon.

In the present study, we explain the data of Ito et al by using a theoretical model of KaiA sequestration^29–31^. This model is based on the large capacity of the KaiC-KaiB-KaiA complexes to absorb KaiA molecules. As illustrated in Figure 1, the KaiC monomer is a tandem array of two homologous domains, the N-terminal domain (CI) and the C-terminal domain (CII)^35^, and these domains are assembled to form the CI ring and the CII ring in a KaiC hexamer^6^. Here we write KaiC hexamer as C_6_. The combined analysis with the cryo-electron microscopy and mass-spectrometry measurements^12^ showed that a KaiB monomer can bind on each CI domain in the CI ring to form KaiC-KaiB complexes, C_6_B_*i*_ with 1 ≤ *i* ≤ 6, and a KaiA dimer further binds on each KaiB to form KaiC-KaiB-KaiA complexes, C_6_B_*i*_A_2*j*_ with 1 ≤ *j* ≤ *i*. This C_6_B_*i*_A_2*j*_ stoichiometry implies the large capacity of KaiC-KaiB-KaiA complexes to absorb significant amount of KaiA molecules. It, therefore, is natural to assume that these KaiC-KaiB-KaiA complexes efficiently sequestrate KaiA. In this study, we use this assumption to explain Ito et al.’s data and propose a practical means to examine the hypothesis of KaiA sequestration.

**Figure 1.**
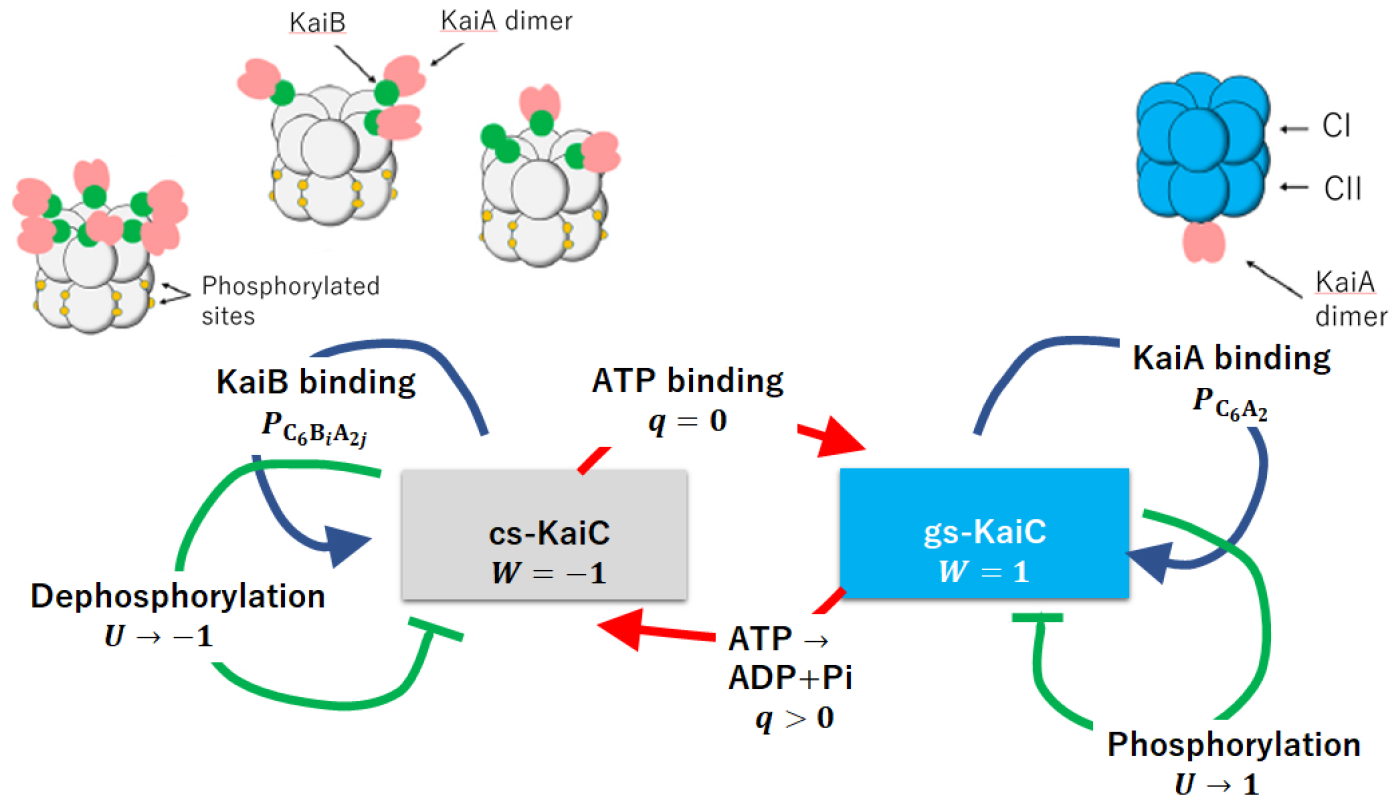
The multifold feedback model of the KaiABC oscillations. The model describes structure transitions, the binding/unbinding of KaiA/KaiB, phosphorylation(P)/dephosphorylation(dP), and ATPase reactions in KaiC hexamers. The KaiC hexamer undergoes structural transitions between the ground state (gs) and the competent state (cs), which are described by a dynamical order parameter *W*(*k, t*); *W*(*k, t*) = −1 when the *k*th KaiC hexamer is in the cs state and *W*(*k, t*) = 1 when it is in the gs state. The probability of binding a KaiA dimer on the CII ring of the *k*th KaiC hexamer is *P*_C_6_A_2__ (*k, t*), and the probability of binding KaiA dimers and KaiB monomers on the CI ring of the kth KaiC hexamer is *P*_C_6_B_i_A_2*j*__ (*k, t*). The phosphorylation level is represented by *U*(*k, t*), where *U*(*k, t*) = −1 when the twelve sites in the CII ring of the *k*th KaiC hexamer are all unphosphorylated and *U*(*k, t*) = 1 when they are all phosphorylated. *q*(*i; k, t*) = 0 when ATP is bound, and *q*(*i; k, t*) = 1 when ADP is bound on the ith CI domain of the kth KaiC hexamer. The KaiA/KaiB binding and the structural transitions establish positive feedback, while the negative feedback works between the P/dP and the structural transitions. Stochastic ATPase reactions trigger the structural transitions. These multifold feedback relations induce switch-like oscillations of individual KaiC hexamers, and those oscillations of different KaiC hexamers are synchronized through the sequestration of KaiA into C_6_B_*i*_A_2*j*_ complexes.

## Multifold feedback model

Our model describes the reactions and structural transitions of KaiC hexamers by considering (i) binding/unbinding of KaiA or KaiB, (ii) phosphorylation (P)/dephosphorylation (dP) in the CII domains, and (iii) ATPase reactions in the CI domains. Because binding/unbinding, P/dP, and ATPase reactions should affect the structure of the KaiC hexamer, and the structure affects the kinetics of those reactions, these structural transitions should constitute multifold feedback (MF) loops. Das, Terada, and Sasai^29–31^ modeled these MF relations in a coarse-grained manner using order parameters of reactions and structure, as illustrated in Figure 1. Here, we consider an ensemble of *N* = 1000 such hexamers and use this MF model for analyzing the experimental data of synchronization of oscillations.

The structural transitions in KaiC hexamer have been observed by the NMR^36,37^, small angle X-ray diffraction^38^, and biochemical^39^ analyses. Each KaiC hexamer undergoes structural transitions between the two states; the state in the P phase and the state in the dP phase. Following Ref. 39, we call the P-phase structure the ground state (gs) and the dP-phase structure the competent state (cs). Here, transitions between the gs and cs states of the *k*th KaiC hexamer are described by a structural order parameter, −1 ≤ *W* (*k, t*) ≤ 1, from *k* = 1 to *N*. *W*(*k, t*) ≈ −1 in the cs state and *W*(*k, t*) ≈ 1 in the gs state at time *t*.

A dimer KaiA can bind on the CII ring of the gs-KaiC, and a monomer KaiB can bind on each CI domain in the CI ring of the cs-KaiC. We describe such dependence of binding affinities on the KaiC structure as

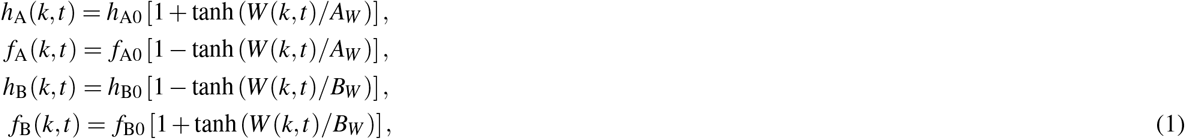

where *h_A_*(*k, t*) and *f*_A_(*k, t*) are the binding and unbinding rate constants of KaiA to and from the CII ring of the *k*th KaiC hexamer at time *t*, respectively, and *h*_B_ (*k, t*) and *f*_B_ (*k, t*) are the binding and unbinding rate constants of KaiB to and from a CI domain of the *k*th KaiC hexamer at time *t*, respectively. Here, *h*_A0_, *f*_A0_, *h*_B0_, and *f*_B0_ are constants to define the time scale, and *A_W_* and *B_W_* are constants to define the structural dependence of the binding affinity.

We write the probability that a KaiA dimer binds on the CII ring of the *k*th KaiC hexamer as P_C_6_A_2__ (*k, t*) and the probability that the number of bound KaiB monomers on the CI ring is *i* and the number of KaiA dimers binding on KaiB is *j* as P_C_6_B_*i*_A_2*j*__ (*k, t*). These probabilities satisfy

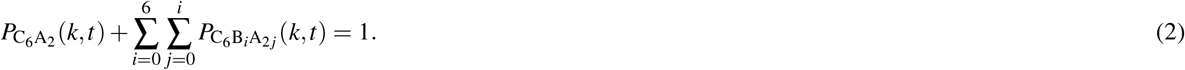

Here, the probability that the kth KaiC hexamer binds neither KaiA nor KaiB at time *t* is P_C_6_B_*i*_A_2*j*__ (*k, t*) with *i = j* = 0.

Each CII domain has two sites to be phosphorylated^26,40^; Ser431, and Thr432. In total, this amounts to twelve sites in a KaiC hexamer. We define −1 ≤ *U* (*k, t*) ≤ 1 to represent the level of phosphorylation in the CII ring of the *k*th KaiC hexamer at time *t* with *U* (*k, t*) = −1 when all the twelve sites are unphosphorylated and *U* (*k, t*) = 1 when they are all phosphorylated. We assume a kinetic equation for *U*(*k, t*) as explained in the Methods section to describe the tendency that *U*(*k, t*) increases when a KaiA dimer binds on the CII ring in the gs state (the P phase), and *U*(*k, t*) decreases when the KaiA dimer unbinds from the CII ring (the dP phase). We assume that the binding of a KaiA dimer on the CII ring stabilizes the gs state and that the binding of KaiB monomers on the CI ring stabilizes the cs state; the latter prevents the KaiA binding on the CII. Furthermore, to ensure the stable oscillation between the two states, we assume that phosphorylation stabilizes the cs state and dephosphorylation stabilizes the gs state. These effects of reactions on the structure can be represented as

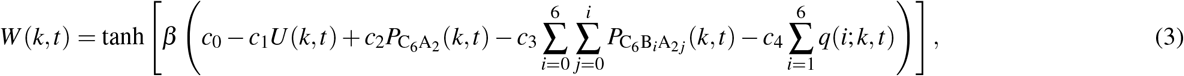

where *β* = 1/(*k*_B_*T*) is the inverse temperature with *k*_B_ being the Boltzmann constant, and *c*_0_, *c*_1_. *c*_2_, *c*_3_, and *c*_4_ are positivevalued constants. Because allosteric transitions in protein oligomers should have a time scale of 10^-3^ ~ 10^-2^ s in general^41^ and the other reactions in the KaiABC system should be much slower than that, we can assume that the structure is in quasi-equilibrium and other chemical states were treated as static constraints to derive the expression of Eq. 3.

We should note that together with Eq. 1, the term *c*_2_P_C_6_A_2__ (*k, t*) in Eq. 3 represents the positive feedback relation between KaiA binding and transition to the gs state. Similarly, the term 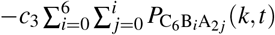 in Eq. 3 represents the positive feedback relation between KaiB binding and transition to the cs state. These positive feedback relations stabilize the gs and cs states, so that the structural changes become switch-like transitions between the two distinct states. The term -*c*_1_*U*(*k, t*) in Eq. 3, on the other hand, represents the negative feedback relation between P/dP reactions and structural transitions. In the gs state with the bound KaiA on the CII ring, the phosphorylation is promoted, which destabilizes the gs state through this term, while in the cs state with the absence of KaiA from the CII ring, the dephosphorylation proceeds, which destabilizes the cs state. Because the time scale of P or dP of twelve sites is ~ 10 hours, the negative feedback effect of P/dP reactions on the structure appears later than the effects of binding/unbinding of KaiA, which takes place in seconds^25^, and binding/unbinding of KaiB, which proceeds in hour^25,28^. Therefore, the negative feedback action through the P/dP reactions should work with some time delay after the structural transition is induced through the positive feedback action of the binding/unbinding reactions. Then, this delayed negative feedback prepares the next structural transition by gradually destabilizing the structural state.

The term 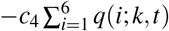 in Eq. 3 represents the effect of ATPase reactions in the CI domains of the *k*th KaiC hexamer at time *t*. KaiC is a slow ATPase enzyme. In the oscillating solution, a KaiC hexamer hydrolyzes ~ 90 ATP molecules/day through the ATPase reactions taking place both in the CI and CII rings^42, 43^. Consumption of ATP molecules in the CII ring supplies phosphate groups to the CII domains for their phosphorylation. Therefore, in the present model, we describe it implicitly with the P/dP kinetics and focus more on the ATPase reactions in the CI. Binding of ATP in the CI is necessary for keeping the hexamer form of KaiC to prevent it from disassembling to monomers^5^. No stable KaiC monomers were observed in the oscillating solution^34^, so we consider that the lifetime of the state in the absence of a bound nucleotide is short enough in the solution with abundant ATP. Therefore, we assume that each CI domain binds either ATP before hydrolysis or ADP after hydrolysis. Thus, we define *q* as

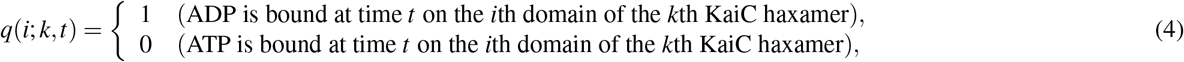

where the ADP release and the subsequent ATP binding are represented by the transition from *q*(*i; k, t*) = 1 to 0, and hydrolysis of the bound ATP is the transition from *q*(*i; k, t*) = 0 to 1. We simulate the stochastic ADP release/ATP binding and hydrolysis by treating *q*(*i; k, t*) as a stochastic variable changing with the lifetime of the ADP bound state, Δ_ADP_, and the frequency of hydrolysis of ATP to ADP + P_i_, *f*_hyd_. Though the role of the ATP hydrolysis in the CI ring is not fully clarified, it was shown that the ATP hydrolysis in the CI is necessary for binding KaiB to KaiC^36,39,44,45^. The binding affinity of KaiB to KaiC is large in the cs state, and thus, it is reasonable to assume that the ATP hydrolysis destabilizes the gs state and promotes transitions to the cs state. This effect is represented in Eq. 3 by the term 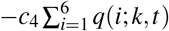. We also assume the positive feedback between ATPase reactions and structural transitions by defining both A_ADP_ and *f*_hyd_ as decreasing functions of *W*(*k, t*). With this definition, the nucleotide state *q*(*i;k, t*) = 1 is stabilized in the cs state, and the nucleotide state *q*(*i;k, t*) = 0 is stabilized in the gs state. In our previous paper^30^, we showed that using this assumption, ATP hydrolysis is a trigger of transition from the gs to cs states, and the ADP release with the subsequent ATP binding is a trigger of transition from the cs to gs states. Thus, the ATPase reactions regulate the timing of structural transitions in the MF model, which explained the experimentally observed correlation between the frequency of oscillations and the rate of ATPase reactions in the CI^29,31^.

Summarizing the above, the structural transitions of KaiC hexamer play a vital role in the oscillations of individual KaiC hexamers in the MF model. The positive feedback between the hexamer structure and the KaiA-CII binding stabilizes the gs state. Similarly, the positive feedback between the hexamer structure and the KaiB-CI binding stabilizes the cs state. After a structural transition of the KaiC hexamer from the cs to gs states or from the gs to cs states, the negative feedback between the hexamer structure and P/dP reactions destabilizes the structural state with some time delay, preparing for the next structural transition. This structural transition is triggered by stochastic ATP binding/hydrolysis.

In the MF model, these oscillations are synchronized through the KaiA sequestration mechanism. We write the total concentration of KaiA on a monomer basis as A_T_ and volume of the solution as *V*. Then, we have the constraint

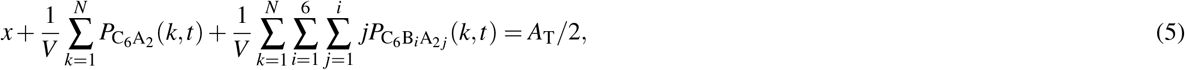

where *x* is the concentration of free unbound KaiA dimer. Because *A*_T_ is a constant of time during oscillations, competition among the three terms in the left-hand side of Eq. 5 couples the dynamics of different KaiC hexamers in the solution; when the third term in the left hand side is large, KaiA is sequestrated into the KaiC-KaiB-KaiA complexes, which decreases the first and second terms in Eq. 5 to suppress the KaiA binding on the CII and reduce the population of the KaiC hexamers in the P phase. In this way, in the present model, the stochastic transitions between the two structural states appear as oscillations in individual KaiC hexamers, and those individual oscillations are synchronized through the sequestration of KaiA into the KaiC-KaiB-KaiA complexes. A further detailed explanation of the model is given in the Methods section.

## Results

We first discuss how the simulated KaiABC system oscillates. We assume that the concentrations of KaiA, KaiB, and KaiC in the system are in the ratio often adopted in in vitro experiments^39,40,42^; A_T_/C_T_ = 1/3 and B_T_/C_T_ = 1, where A_T_, B_T_, and C_T_ are concentrations of KaiA, KaiB, and KaiC on a monomer basis, respectively, with C_T_ = 6*N/V*. For the ease of comparing the simulated results with the data in the experimental reports, we transform the theoretical variables *U* and *W* ranging from −1 to 1 to the variables *D* and *X* ranging from 0 to 1 and define their ensemble averages 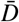 and 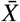 as

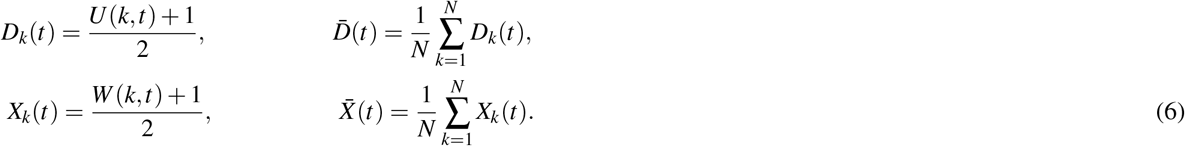

In Figure 2a, we show the oscillations of the phosphorylation level *D_k_* (*t*) and the structural state *X_k_*(*t*) of a single KaiC hexamer, which was arbitrarily chosen from the simulated ensemble of *N* = 1000 KaiC hexamers. Figure 2a shows that the single molecule undergoes switch-like transitions between the cs (*X_k_* ≈ 0) and gs (*X_k_* ≈ 1) states. These discrete transitions arise from the positive feedback relations among reactions and structural changes in the KaiC hexamer. On the other hand, the phosphorylation level *D_k_* follows these cooperative switching transitions with a slower rate to show the oscillations between P and dP processes. Figure 2b shows the oscillations of the ensemble, in which the fluctuations in the single-molecular stochastic oscillations are averaged, leading to the smoothed coherent oscillations of the phosphorylation level 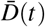 and the structural state 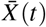. Also plotted in Figure 2 are the nucleotide bound state *q_k_* (*t*) in the CI domains of a single molecule and 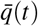 in the ensemble, which are defined as

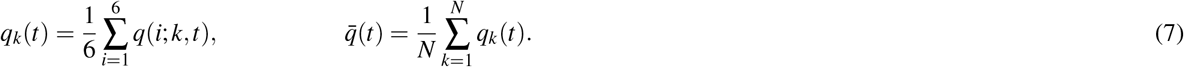

**Figure 2.**
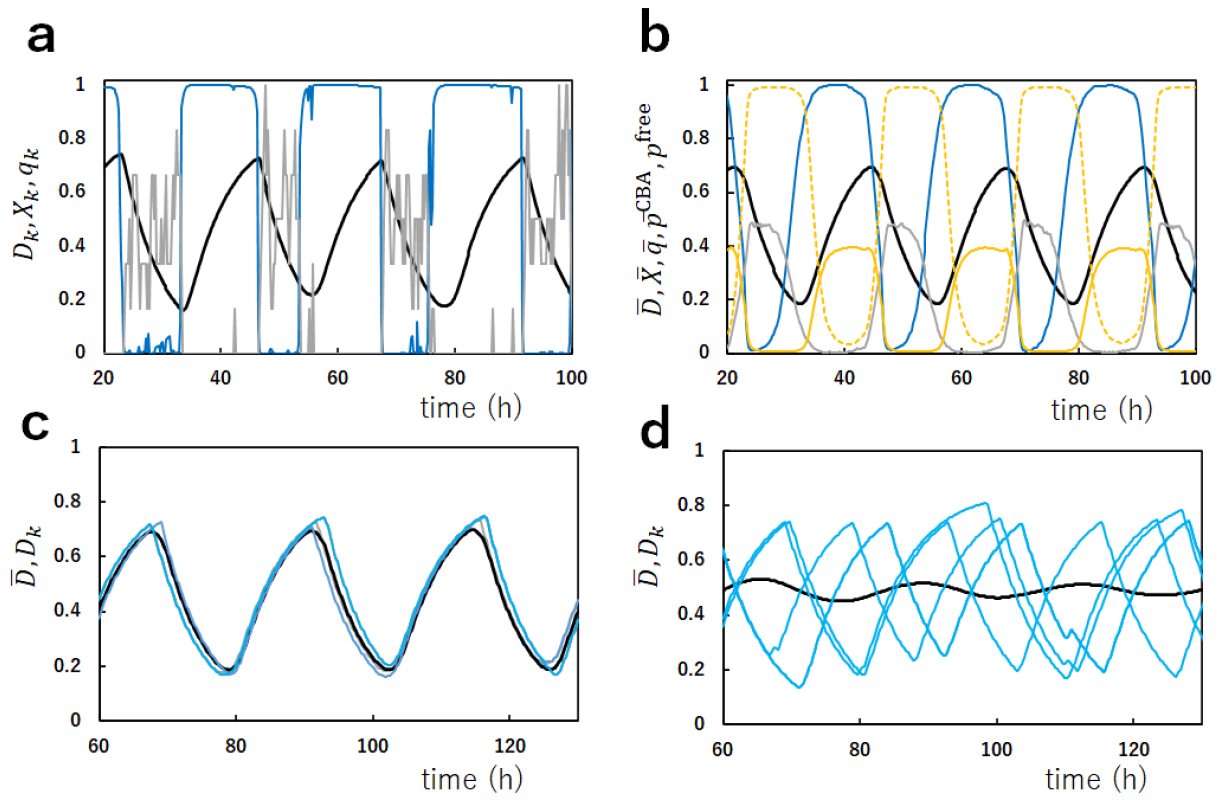
Single-molecule and ensemble-level oscillations of the simulated KaiABC system. (**a**) Oscillations of an example single KaiC hexamer arbitrarily chosen from the ensemble of *N* = 1000 KaiC hexamers. The phosphorylation level *D_k_*(*t*) (black), the structural state *X_k_*(*t*) (cyan), and the nucleotide state of the CI ring *q_k_*(*t*) (gray) are shown. (**b**) Oscillations of the ensemble of *N* = 1000 KaiC hexamers. The phosphorylation level 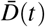 (black), the structural state 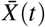 (cyan), the nucleotide state of the CI ring 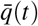 (gray), the normalized concentration of free KaiA dimers *p*^free^(*t*) (orange), and the fraction of the sequestrated KaiA dimers 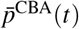 (orange dashed line) are shown. (**c, d**) The ensemble oscillations of the phosphorylation level 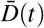 (black) and oscillations of the phosphorylation level *D_k_*(*t*) (cyan) of five arbitrarily chosen examples of single KaiC hexamers are superposed. The dissociation constant between the KaiC-KaiB complexes and KaiA is 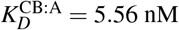 in **a, b** and **c**, and 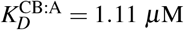 in **d**.

In the single-molecular oscillations in Figure 2a, *q_k_*(*t*) shows distinct switching between the ATP bound (*q_k_*(*t*) ≈ 0) and ADP bound (*q_k_*(*t*) ≈ 0.5) states. The nucleotide bound state in the six CI domains in a hexamer shows cooperative switching, which arises from the positive feedback between ATPase reactions and structural changes. Figure 2b shows that the nucleotide state at the ensemble level 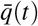 shows smooth oscillations, which have peaks in the dP phase.

The fraction of the KaiA molecules sequestrated into the KaiC-KaiB-KaiA complexes and the normalized concentration of free unbound KaiA dimer are plotted in Figure 2b by defining

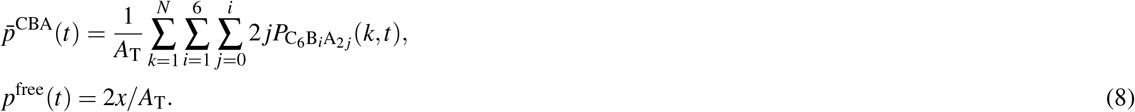

The fraction of the sequestrated KaiA, 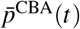, is small and the concentration of free unbound KaiA, *p*^free^(*t*), is finite in the P phase, while 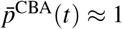 and *p*^free^(*t*) ≈ 0 in the dP phase, showing that KaiA is available for binding the CII ring in the P phase but is sequestrated and not available to the CII in the dP phase. This sequestration is the mechanism of synchronization in the present MF model. In the previous papers, we showed with the MF model^29,31^ that the synchronization due to this KaiA sequestration is lost when the amount of KaiA is too large in the simulated ensemble, diminishing the amplitude of the ensemble oscillations. The disappearance of the ensemble oscillations in our simulations agrees with the experimental data^33^, supporting the KaiA sequestration hypothesis assumed in the present model. A similar effect is seen when the binding affinity of KaiA to the KaiC-KaiB complexes is reduced by increasing the dissociation constant 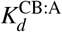 in the model. In Figure 2c the ensemble oscillations, 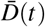, and the single-molecular oscillations, *D_k_*(*t*), of arbitrarily chosen five KaiC hexamers are superposed, showing that individual oscillations are synchronized to generate the coherent ensemble oscillations. This synchronization is lost in Figure 2d when 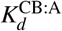 is increased, showing the importance of the sequestration mechanism for the synchronization.

The synchronization mechanism can be further analyzed by simulating the mixture solution. Figure 3 shows examples of the simulated mixing. From two oscillating ensembles, *e*_1_ and *e*_2_, each of which consists of *N* KaiC hexamers, *N*/2 hexamers were arbitrarily chosen and mixed, resulting in the mixture ensemble of *N* hexamers. Figure 3 shows that when *e*_1_ at the lowest phosphorylation level and *e*_2_ at the highest phosphorylation level are mixed, the resultant mixture ensemble is entrained into the oscillations starting from the highest phosphorylation level (Figure 3a) and when *e*_1_ in the dP phase and *e*_2_ in the P phase are mixed, the resultant mixture ensemble is entrained into the dP phase (Figure 3b).

**Figure 3.**
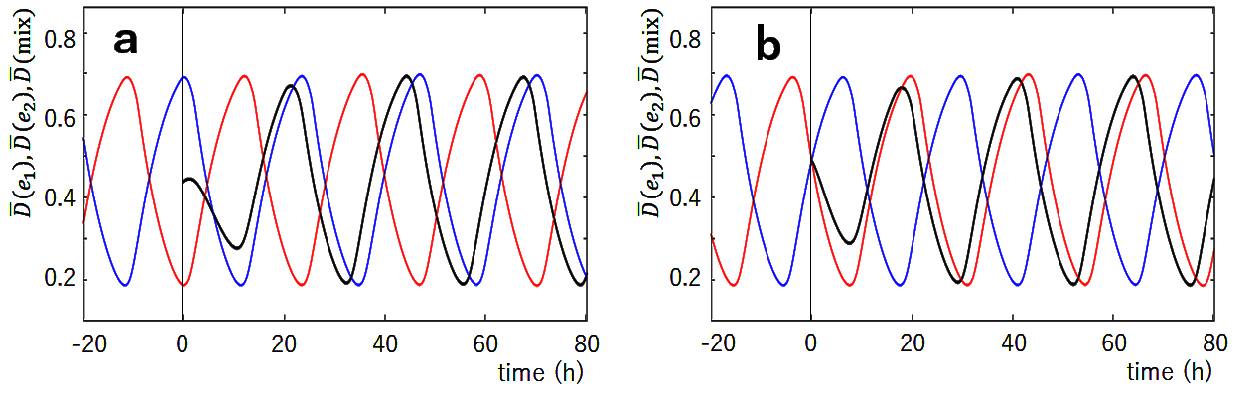
The simulated mixing of two ensembles, *e*_1_ and *e*_2_, oscillating in different phases. The phosphorylation level of *e*_1_, 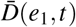 (red), and the phosphorylation level of *e*_2_, 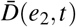 (blue), are superposed with the phosphorylation level after mixing, 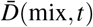 (black). *e*_1_ and *e*_2_ were mixed at *t* = 0 designated by a vertical line. (**a**) *e*_1_ at the lowest phosphorylation level and *e*_2_ at the highest phosphorylation level were mixed. (**c**) *e*_1_ in the dephosphorylation phase and *e*_2_ in the phosphorylation phase were mixed. Parameters are the same as in Figures 2a–2c.

These entrainment features can be examined more extensively by choosing six different phases from A to F, as shown in Figure 4a. In Figure 4b, we mixed the equivalent amount of ensembles, *e*_1_ and *e*_2_, and plotted the time when the peaks of oscillations in the ensemble appear after the mixing. In each panel of Figure 4b, the phase of *e*_1_ was fixed and the phase of *e*_2_ was varied from A to F. When oscillations of the mixture ensemble are entrained into the oscillations in the phase of *e*_1_, then the peaks are aligned along the vertical line. On the other hand, when oscillations of the mixture ensemble are entrained into the oscillations in the phase of *e*_2_, then the peaks are aligned near the slant line. Comparing with Figure 2c, we found the simulated results well explain the data of Ito et al.^13^; oscillations of the mixture solution are entrained into the oscillations in the phases of the dP process, C, D, and E in Figure 4a. This agreement of the simulated results with the experimental observations supports the hypothesis that the KaiA sequestration into the KaiC-KaiB-KaiA complexes explains the essential part of the mechanism of synchronization. The present results showed that the monomer exchange or the KaiA sequestration into the lowly phosphorylated states in the P phase or in the dP phase is not necessary for explaining Ito et al.’s experimental data on entrainment.

**Figure 4.**
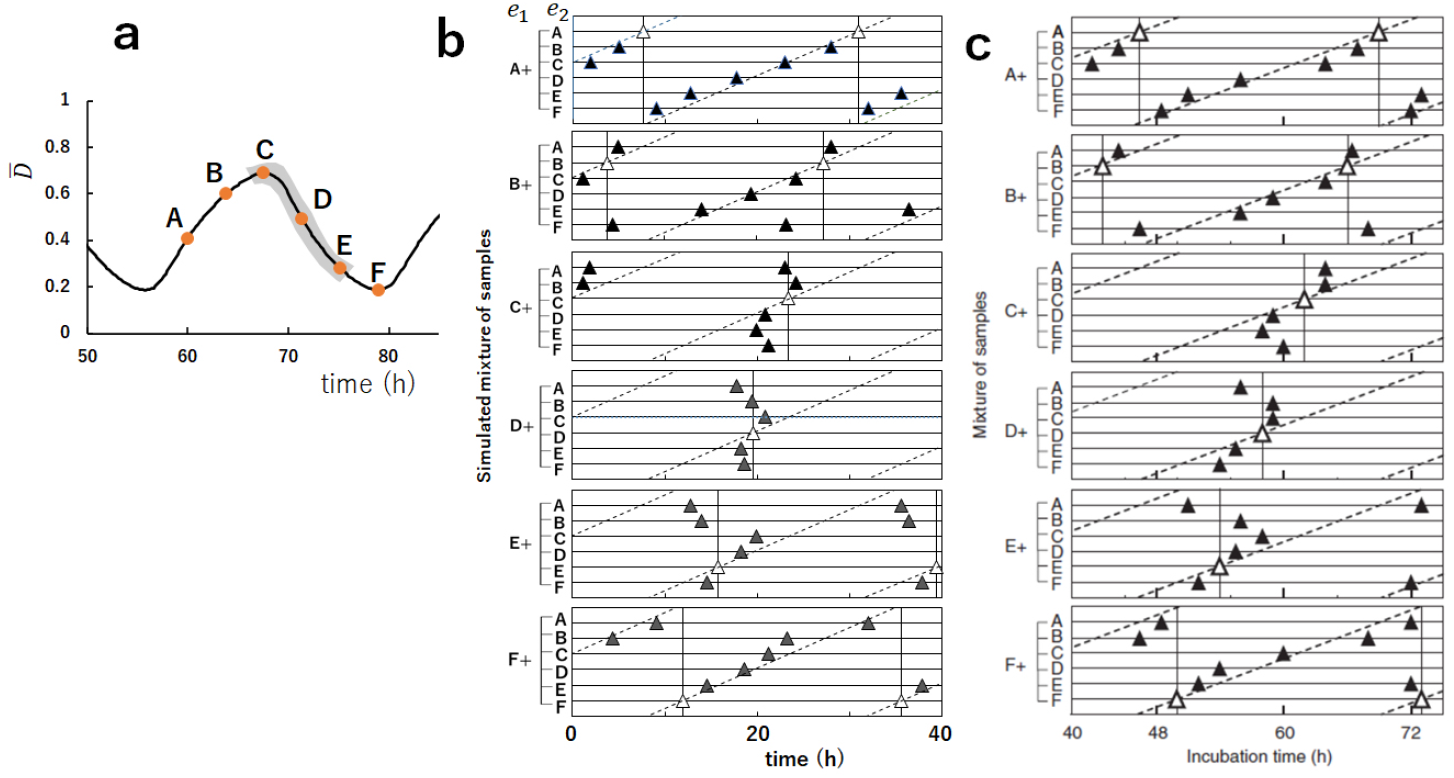
The simulated mixing of two ensembles oscillating in different phases from A to F. (**a**) Six phases, A to F, were selected from the simulated curve of the phosphorylation level 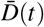. The simulated results show that the phase of the oscillations of the mixture ensemble are entrained into the phase of gray shaded region. (**b**) Time when the peaks of 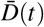 appeared after mixing two samples, *e*_1_ and *e*_2_, is plotted by triangles. White triangles when the same samples were mixed, and filled triangles when samples in different phases were mixed. In each of six panels, the phase of *e*_1_ was fixed and the phase of *e*_2_ was varied from A to F. In **a** and **b**, parameters are the same as in Figure 3. (**c**) The plot of the experimental data corresponding to **b**. Copied from Ref.13 with permission.

The present model suggests that the efficient KaiA sequestration into the KaiC-KaiB-KaiA complexes in the dP phase should be the mechanism of entrainment of oscillations into the dP phase. When the ensemble in the dP phase and the ensemble in the P phase are mixed, the free unbound KaiA dimers supplied from the P-phase ensemble are absorbed by the KaiC-KaiB-KaiA complexes supplied from the dP-phase ensemble. Then, the KaiC hexamers in the gs state in the mixture ensemble are turned into the cs state because of the lack of free KaiA, which can stabilize the gs state by binding to the CII, which further increases the population of the KaiC-KaiB-KaiA complexes. This scenario of entrainment can be tested by comparing the predictions of simulations with the experimental measurements. In Figure 5, mixing of the solution at the highest phosphorylation level and the solution at the lowest phosphorylation level was simulated with different concentrations of KaiB or KaiA to show how the modification of the efficiency of the KaiA sequestration affects the synchronization. The sequestration efficiency is decreased when the concentration of KaiB, *B*_T_, is decreased, which, in turn, decreases *P*_C_6_B_*i*_A_2*j*__ (Figure 5a), or when the concentration of KaiA, *A*_T_, is increased, which, in turn, weakens the effect of constraint Eq. 5 (Figure 5b). Figure 5a shows that with the larger *B*_T_, the entrainment into the Phase-C oscillation is not changed, while with the smaller *B*_T_, the entrainment becomes obscure. It is intriguing that with a moderately small *B*_T_ as *B*_T_/*C*_T_ = 4/6, oscillations are still coherent with the large amplitude in a stable condition, but oscillations are weakened upon mixing because of the weak entrainment tendency and the recovery of the amplitude is slow after the mixing. By further decreasing *B*_T_, the synchronization is lost and the ensemble oscillations disappear at *B*_T_/*C*_T_ ≈ 0.5. As shown in Figure 5b, the results of the change in *A*_T_ are more complex. With the larger *A*_T_, the entrainment becomes weaker, and the oscillation disappears for *A*_T_/*C*_T_ > 1. With the smaller *A*_T_, as in the case of *A*_T_/*C*_T_ = 1/6, the entrainment tendency is not altered from the standard case of *A*_T_/*C*_T_ = 1/3, which is because the constraint Eq. 5 works efficiently for the smaller *A*_T_. However, with the further smaller *A*_T_, the entrainment becomes weaker because of the smaller *P*_C_6_B_*i*_A_2*j*__. These simulated results are testable in the experiment by changing the concentration of KaiA or KaiB.

**Figure 5.**
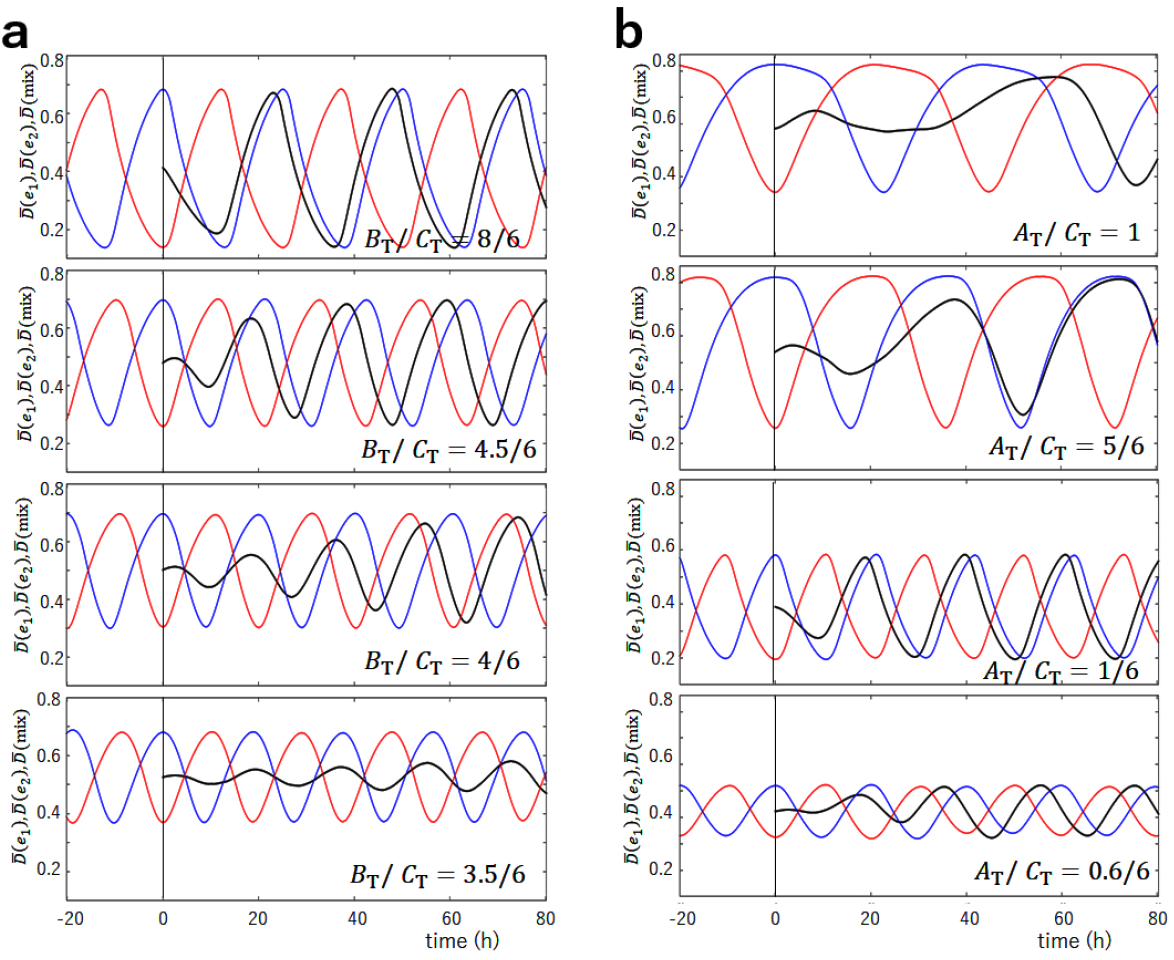
The simulated mixing of two ensembles, *e*_1_ at the lowest phosphorylation level and *e*_2_ at the highest phosphorylation level. The phosphorylation level of *e*_1_, 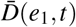 (red), and the phosphorylation level of *e*_2_, 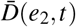 (blue), are superposed with the phosphorylation level after mixing, 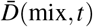 (black). *e*_1_ and *e*_2_ were mixed at *t* = 0 designated by a vertical line. (**a**) The ratio of the concentration of KaiB over the concentration of KaiC on a monomer basis, *B*_T_/*C*_T_, is varied from the standard value of *B*_T_/*C*_T_ = 1. (**b**) The ratio of the concentration of KaiA over the concentration of KaiC on a monomer basis, *A*_T_/*C*_T_, is varied from the standard value of *A*_T_/*C*_T_ = 2/6. Other parameters are the same as in Figure 3.

## Discussion

The MF model, which considers the multifold feedback relations among binding/unbinding of KaiA or KaiB, phosphorylation/dephosphorylation, ATPase reactions, and the KaiC structural transitions, predicts the switch-like transitions of structure and ATPase activity of individual KaiC hexamers. As KaiA is sequestrated into the KaiC-KaiB-KaiA complexes, these switch-like transitions in individual molecules are synchronized to give rise to the coherent oscillations in the ensemble. In this study, we showed that this model quantitatively reproduced the experimental data on entrainment in the mixture solution, supporting the validity of the mechanism of KaiA sequestration into the KaiC-KaiB-KaiA complexes. The present results calculated without assuming the monomer exchange suggest that the KaiA sequestration should be a dominant mechanism of synchronization though the monomer-exchange mechanism and the KaiA sequestration mechanism are not mutually exclusive. We showed that this hypothesis could be examined experimentally through the biochemical assays by varying concentrations of KaiA or KaiB.

Finally, we should emphasize the role of ATPase reactions in the CI domains in synchronization. A previous paper^31^ shows that synchronization is lost in the MF model when the frequency of ATPase reactions decreases. With the lower frequency of ATPase reactions, the frequency of structural transitions in individual KaiC hexamers decreases, which enhances the structural fluctuation in KaiC hexamer, and ultimately diminishes the effects of synchronization necessary for the coherent ensemble oscillations. This is an example that the free energy of ATP is consumed for communicating among molecules. The free energy consumption necessary for molecular communication was also discussed by assuming the monomer-exchange mechanism with a simplified model of KaiC oscillations^20^. Further quantitative analyses on the ATP consumption in the KaiABC system could lead to a deeper understanding of the thermodynamics of molecular communication. It is intriguing to examine whether a similar mechanism based on the coupling between the molecular sequestration and the ATP-induced transitions in the binding affinity in the other biological regulatory systems. Further analyses of ATPase reactions and synchronization should help develop the framework for understanding molecular communication.

## Methods

### Binding/unbinding reactions of KaiA and KaiB

The master equation for the probability distribution of chemical states should represent the kinetics of stochastic reactions. When the probability distribution of the many-body system is approximately factorized into the single-molecular probability distributions as *P*_C_6_A_2__ (*k, t*) and others, the master equation can be reduced to simpler equations similar to the chemical kinetics equations^46,47^. It follows that the equation for *P*_C_6_A_2__ (*k, t*) should be 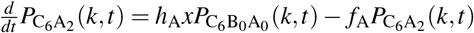. However, the recent observations with atomic force microscopy showed that the KaiA binding/unbinding process has a time scale of seconds^25^, which is much smaller than the time scales of KaiB binding/unbinding and P/dP. Therefore, we consider that binding/unbinding of KaiA is in quasi-equilibrium as 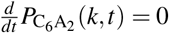. Then, we have

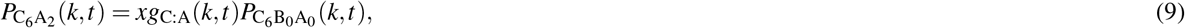

where *g*_C:A_(*k,t*) = *h*_A_(*k,t*)/*f*_A_(*k,t*). Similarly, binding/unbinding of KaiA to KaiB should be in quasi-equilibrium, leading to

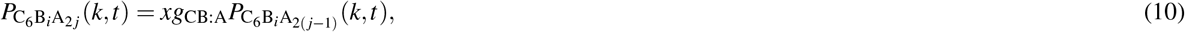

where *g*_CB:A_ is a constant representing the binding affinity of KaiA to the KaiC-KaiB complexes. Because KaiA does not directly interact with KaiC in this binding/unbinding to/from KaiB, we assume that *g*_CB:A_ does not depend on the KaiC structure *W*(*k, t*). With this expression, we can write 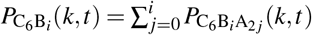 and 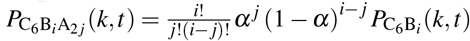 with 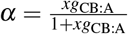. Then, the constraint Eq. 5 is

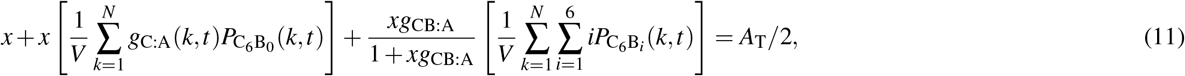

from which *x* at time *t* is determined. The equations for the binding/unbinding of KaiB are

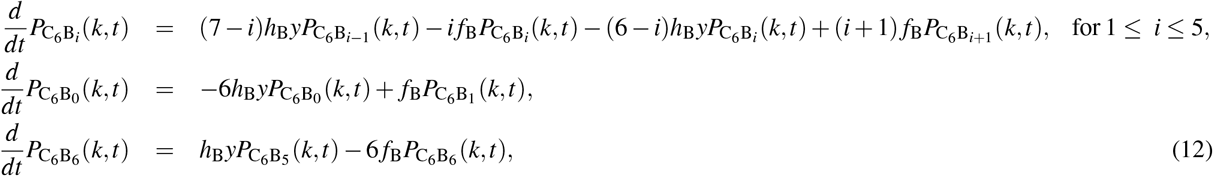

where *y* is the concentration of free unbound KaiB monomers. The constraint coming from the conservation of total number of KaiB molecules is

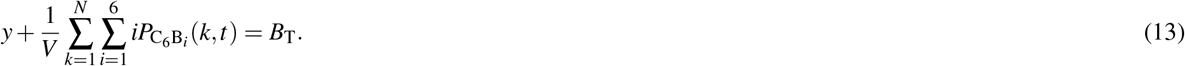

### Phosphorylation/dephosphorylation reactions

We assume that the kinetics of phosphorylation and dephosphorylation depend on whether a KaiA dimer binds on the CII as

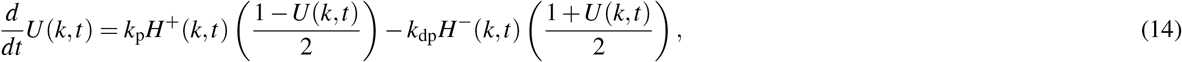

where *H*^+^ (*k, t*) = *z*/(1 + *z*) and *H* (*k, t*) = 1/ (1 + *z*) represent the effects of binding and unbinding of KaiA to and from the CII, respectively, with *z* = P_C_6_A_2__ (*k, t*)/*P*_0_. Here, *k*_p_, *k*_dp_, and *P*_0_ are constants.

### ATPase reactions

We assume that the bound ATP on the ith CI domain of the *k*th KaiC hexamer is hydrolyzed into ADP and P_i_ at a random timing with the frequency

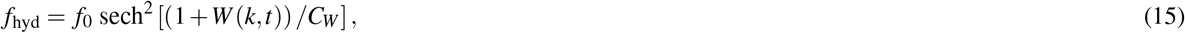

with constants *f*_0_ and *C_W_*. With this frequency, the nucleotide bound state of the ith domain is changed from *q*(*i; k, t*) = 0 to 1. It is not certain whether this hydrolysis immediately impacts the structure to stabilize the cs state or the effect is significant only after the P_i_ is released. We do not distinguish these two cases in the present mathematical formulation, but we write the nucleotide state that stabilizes the cs state as *q*(*i; k, t*) = 1, which is either the ADP+P_i_ bound state or the state in which ADP remains bound after the release of P_i_. We assume that ADP is kept bound at the ith domain of the *k*th hexamer for the time duration 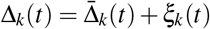, where *ξ_k_*(*t*) is a random number satisfying 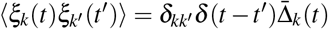. The average lifetime of the ADP bound state 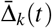 depends on the structure *W* (*k, t*) as

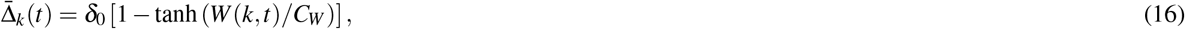

where *δ*_0_ is a constant. After the ADP is released, the next ATP binds to the same site, which turns the nucleotide state from *q*(*i; k, t*) = 1 to 0.

### Parameters

We simulated the system which contains *N* = 1000 KaiC hexamers at temperature *T* = *T*_0_ = 30°C. For *N* = 1000 and *V* = 3 × 10^-15^*l*, the concentration of KaiC is *C*_T_ = 3.3μM on a monomer basis, which is close to the 3.5μM concentration often used in experiments. We assume the ratio *A*_T_: *B*_T_: *C*_T_ = 1: 3: 3 as adopted in many experiments^39,40,42^ except for the results shown in Figure 5. The oscillations are robust against small changes in the rate constants; therefore, the parameters of the rate constants were not finely tuned but determined from the order of magnitude argument.

In units of *V* = 1, the binding rate constant *h*_B0_*B*_T_ and the unbinding rate constant *f*_B0_ between KaiB and the CI were chosen to be *h*_B0_*B*_T_ ≈ *f*_B0_ ≈ 1 h^-1^. Here, we used *h*_B0_ = (5/6) × 10^-4^ h^-1^ and *f*_B0_ = 1 h^-1^, corresponding to the dissociation constant of 6.67 μM for *V* = 3 × 10^-15^*l*, which is further modulated by the structural change of KaiC as in Eq. 1. The dissociation constant of binding between KaiA and the CII was set to be 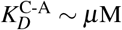 as observed experimentally^34^. We used *h*_A0_/*f*_A0_ = 5 × 10^-4^ in units of *V* = 1, corresponding to 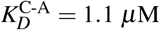 for *V* = 3 × 10^-15^*l*, which is further modulated by the structural change as in Eq. 1. The dissociation constant of KaiA and the KaiC-KaiB complexes was not yet observed experimentally. Here, we assumed a rather small dissociation constant to ensure the sequestration effect. We used *g*_CB:A_ = 1 × 10^-1^ in units of *V* = 1, which corresponds to 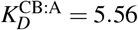 nM for *V* = 3 × 10^-15^*l*, except for the results shown in Figure 2d. 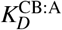 does not depend on the structure of KaiC in the present model.

The constants for the P/dP reactions were chosen to be *k*_p_ = *k*_dp_ = 3 × 10^-1^ h^-1^, and *P*_0_ = 0.1. The constants for the ATPase reactions were *f*_0_ = 1 h^-1^ and *δ*_0_ = 1 h. The constants representing the coupling between reactions and structure were *c*_0_ = 2*k*_B_T_0_, *c*_1_ = *c*_2_ = *c*_3_ = 4*k*_B_T_0_, and *c*_4_ = *k*_B_T_0_. The constants defining the dependence of the rates on the structure were *A_W_* = *B_W_* = *C_W_* = 1.

## Acknowledgements

This work was supported by JST-CREST Grant JPMJCR15G2, the Riken Pioneering Project, and JSPS-KAKENHI Grants JP19H01860, 19H05258 and 20H05530.

